# PSL-Recommender: Protein Subcellular Localization Prediction using Recommender System

**DOI:** 10.1101/462812

**Authors:** Ruhollah Jamali, Changiz Eslahchi, Soheil Jahangiri-Tazehkand

## Abstract

Identifying a protein’s subcellular location is of great interest for understanding its function and behavior within the cell. In the last decade, many computational approaches have been proposed as a surrogate for expensive and inefficient wet-lab methods that are used for protein subcellular localization. Yet, there is still much room for improving the prediction accuracy of these methods.

PSL-Recommender (Protein subcellular location recommender) is a method that employs neighborhood regularized logistic matrix factorization to build a recommender system for protein subcellular localization. The effectiveness of PSL-Recommender method is benchmarked on one human and three animals datasets. The results indicate that the PSL-Recommender significantly outperforms state-of-the-art methods, improving the previous best method up to 31% in F1 – mean, up to 28% in ACC, and up to 47% in AVG. The source of datasets and codes are available at: https://github.com/RJamali/PSL-Recommender

## 1. Introduction

Proteins are responsible for a wide range of functions within cells. The functionality of a protein is entangled with its subcellular location. Therefore, identifying Protein Subcellular Localization (PSL) is of great importance for both biologists and pharmacists, helping them inferring a protein’s function and identifying drug-target interactions [1]. Recent advances in genomics and proteomics provide massive amount of protein sequence data extending the gap between sequence and annotation data. Although PSLs can be identified by experimental methods, these methods are laborious and time-consuming explaining why only a narrow range of PSL information in Swiss-Prot database has been verified in this manner [2]. This problem augments the demand for accurate computational prediction methods. Developments of computational and machine learning techniques have provided fast and effective methods for PSL prediction [2–5, 5–23].

The desired PSL prediction can be reached typically by relying on sequence-derived features, taking into consideration that using annotation-derived features can lead up to better performance. Different types of sequence-derived features have been used for PSL prediction. For example, PSORT [24], WoLF PSORT [4] and TargetP [25] employ sequence sorting signals [9] while Cell-Ploc [10] and LOCSVMPSI [11] use position specific scoring matrix [26]. Additionally, amino/psudo-amino acid composition information [12, 27] is utilized by ngLOC [13]. There are also some methods that employ combinations of sequence based features [3, 4]. Alongside, there are different types of annotation derived features such as protein-protein interaction, Gene Ontology (GO) terms and functional domain and motifs which are used by different methods [2, 7, 8, 17–20]. Moreover, text-based features derived by literature mining have also been employed beside other features for protein subcellular localization [14–16].

Parallel to the importance of features, selecting a suitable algorithm definitely leads to a higher accuracy in prediction. Many machine learning methods or statistical inferences are applied for the protein subcellular localization problem, such as support vector machine [2, 3], K-nearest neighbors [21, 22], and Bayesian methods [6, 23].

In this paper, we have modeled the PSL prediction problem as a recommendation task that aims to suggest a list of subcellular locations to a new protein. In general, Recommendation systems are methods and techniques that suggest users a preferred list of items (e.g. suggesting a movie to watch or suggesting an item to purchase) based on a previous knowledge about relations within and between items and users [28]. As of late framework strategies, recommendation systems have been utilized to predict associations in challenging bioinformatics problems [29–32]. Well-known PSL prediction methods assign equal importance to all proteins information in both constructing model and prediction tasks [2, 5, 6, 33], but utilization prioritized information from similar proteins in model construction step is likely more meaningful. Additionally, due to large number of protein features, dimension reduction methods which capture dependencies among proteins and subcellular locations could be useful to construct a PSL prediction model. In order to considering these concepts, our method, “PSL-Recommender” employs a probabilistic recommender system to predict the presence probability of a protein in a subcellular location. PSL-Recommender utilized both prioritized information to elucidate the importance of sharing similarity information over proteins, and low-dimensional latent space projection of protein features during PSL prediction process.

PSL-Recommender employs logistic matrix factorization technique [34] integrated with a neighborhood regularization method to capture the information from a set of previously known protein-subcellular location relations. Then, it utilizes this information to predict the presence probability of a new protein in a subcellular location using a logistic function. Logistic Matrix factorization has shown promising results for problems such as music recommendation [35], drug-target interaction prediction [29, 36], and lncRNA-protein interaction prediction [30]. However, to the best of our knowledge, it has not been used in PSL prediction problem.

By evaluating on different benchmark datasets, we have shown that PSL-Recommender significantly outperforms the results of current state-of-art methods.

## 2. Materials and Method

### 2.1. Method

To recommend a subcellular position to a protein, PSL-Recommender employs two matrices; a matrix of currently known protein-subcellular location assignments(PSL interactions) and a similarity matrix between proteins. The proteins similarity matrix is the weighted average of similarity measures such as GO terms [37] similarities, PSSM [38] similarity and STRING [39] similarity. The main idea is to model the localization probability of a protein in a location as a logistic function of two latent matrices. The latent matrices are acquired by matrix factorization of the protein-subcellular location matrix with respect to the similarity matrices. Construction pipeline of PSL-Recommender predictor has been demonstrated in Fig 1. The details of similarity measures and the recommender system are as follows.

**Figure 1:**
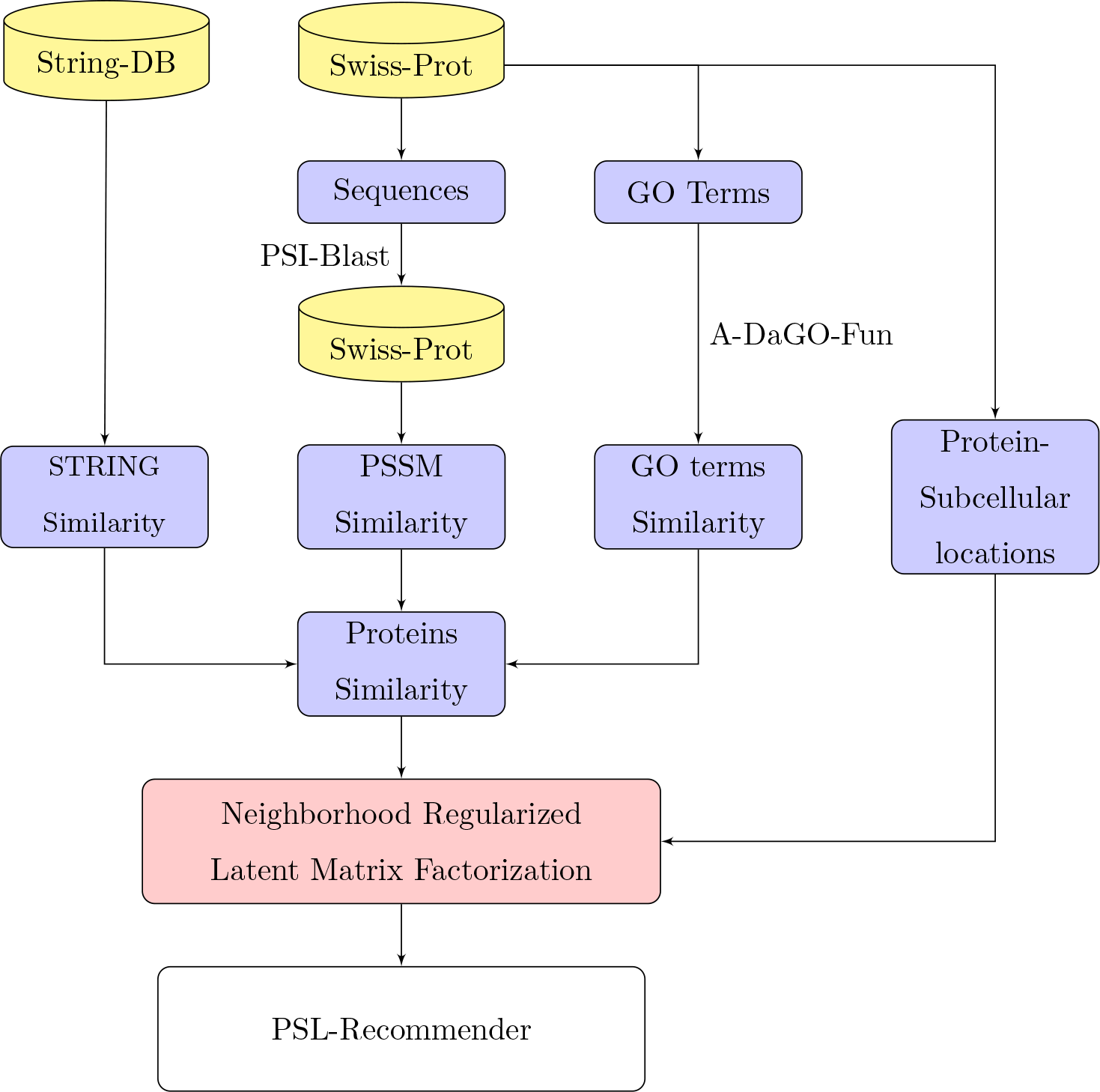
Construction pipeline of PSL-Recommender

#### 2.1.1. PSSM similarity

The PSSM similarity matrix, 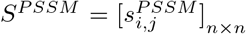, contains the pairwise global alignment scores of proteins that are calculated using the position specific scoring matrices (PSSM). Accordingly, to compute the 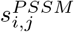 of proteins *i* and *j*, first for each protein, PSI-BLAST [40] with e-value 0.001 is used to search the Swiss-Prot database to obtain each protein’s PSSM. Then *i* and *j* are globally aligned twice, once using the PSSM of *i* and once using the PSSM of *j*. Finally, 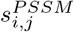 is obtained by the mean of reciprocal alignment scores. The PSSM similarity matrix is normalized using unity based normalization.

#### 2.1.2. STRING similarity

It has been shown that two interacting proteins have a higher chance to be in the same subcellular location [8, 18, 41]. Accordingly, we extracted the interaction score of all pairs of proteins from STRING (Ver. 10.5) to construct the proteins interaction scoring matrix. If no interaction was available for a pair of proteins, we set their interaction score to zero. Since the STRING protein-protein interaction scores are in the range of [0, 999], we normalized the scores with unity-base normalization.

#### 2.1.3. Semantic similarity of GO terms

Gene Ontology terms are valuable sources of information for predicting subcellular localization [2, 42]. To exploit GO terms similarities, we first extracted GO molecular function, biological process and cellular component terms from Swiss-Prot database. Then we used A-DaGO-Fun to extract the BMA-based Resnik GO terms semantic similarities [43]. Similarities were normalized using unity-based normalization.

#### 2.1.4. PSL-Recommender

Let proteins and subcellular locations sets be denoted by X and Y, respectively and |*X*| = *m* and |*Y*| = *n*. Moreover, let 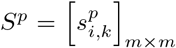 represent the similarity of proteins. The presence of proteins in subcellular locations is also denoted by a binary matrix *L* = [*l*_*ij*_]_*m*×*n*_, where, *l*_*ij*_ = 1 if proteins *i* has been experimentally observed in subcellular location *j* and *l*_*ij*_ = 0 otherwise.

The localization probability of the protein *i* in subcellular location *j* can be modeled as a logistic function as follows:

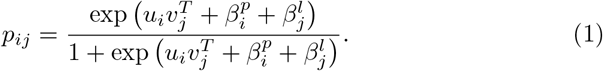

In Eq.(1), 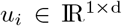 and 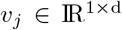 are two latent vectors that reflect the properties of protein *i* and subcellular location *j* in a shared latent space of size *d* < *min* (*m*, *n*). However, in our case matrix *L* is biased toward some proteins and subcellular locations, meaning that some proteins tend to localize in many locations and some subcellular locations include many proteins. Accordingly, for each protein and subcellular location we introduce a latent term to capture this bias. In Eq.(1), 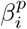 represent the bias factor for protein *i* and 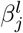 represent the bias factor for subcellular location *j*.

Now the goal is to acquire the latent factors for a given *L*. Suppose 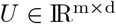, 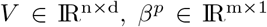 and 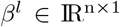 denote the latent matrices and bias vectors for proteins and subcellular locations. According to the Bayes’ theorem and the independence of *U* and *V* we have:

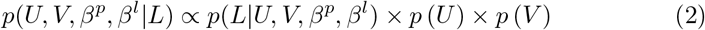

On the other hand, by assuming that all entries of *L* are independent, we have:

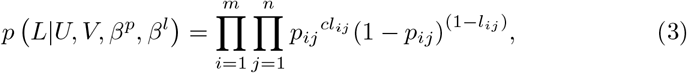

where *c* is weighting factor on positive observations, since we have more confidence on positive observations than negative ones. Also, by placing a zero-mean spherical Gaussian prior on latent vectors of proteins and subcellular locations we have:

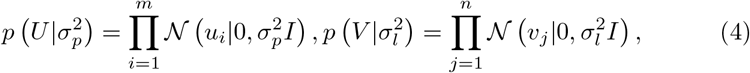

where 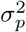 and 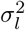 are parameters controlling the variances of prior distributions and *I* denotes the identity matrix. According to the above equations, the log of the posterior is yielded as follows:

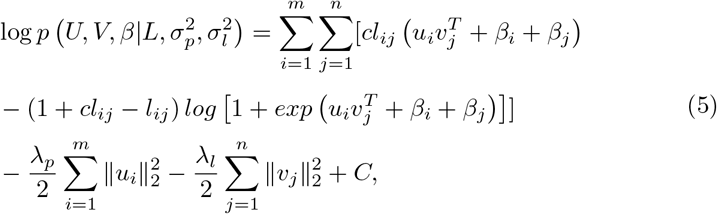

where 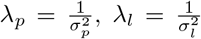 and *c* is a constant term independent of the model parameters. Our goal is to learn *U*, *V*, *β*^*p*^ and *β*^*l*^ that maximize the log posterior above, which is equal to minimizing the following objective function:

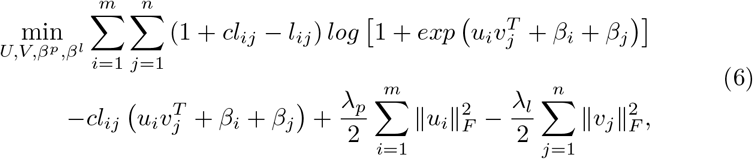

where ‖.‖_*F*_ denotes the Frobenius norm of a matrix. By minimizing the above function *U*, *V* and *β* can effectively capture the information of protein localizations. However, we can further improve the model by incorporating protein similarities as suggested by [29]. This process is known as neighborhood regularization. This is done by regularizing the latent vectors of proteins such that the distance between a protein and its similar proteins is minimized in the latent space.

Accordingly, suppose that the set of *k*_1_ most similar neighbors to protein *x*_*i*_ is denoted by 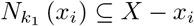. We constructed adjacency matrix *A* = [*a*_*ij*_]_*m*×*m*_ that represents proteins neighborhood information as follows:

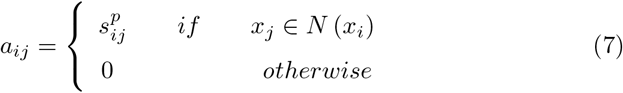

To minimize the distance between proteins and their *k* most similar proteins we minimize the following objective function:

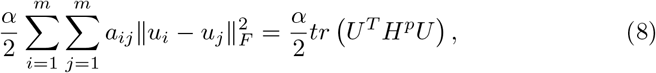

where 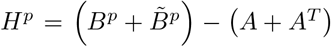 and *tr* (.) is the trace of matrix. In this equation, *B*^*p*^ and 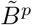 are two diagonal matrices, that their diagonal elements are 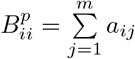 and 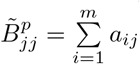, respectively.

Finally by plugging Eq.(8) into Eq.(6) we will have the following:

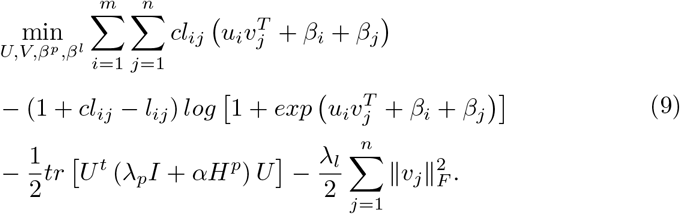

A local minimum of above function can be found by employing the alternating gradient descent method. In each iteration of the gradient descent, first *U* and *β*_*i*_ are fixed to compute *V* and *β*_*j*_ and then *V* and *β*_*j*_ are fixed to compute *U* and *β*_*i*_. To accelerate the convergence, we have employed the AdaGrad [44] algorithm to choose the gradient step size in each iteration adaptively. The partial gradients of latent vectors and biases are given by:

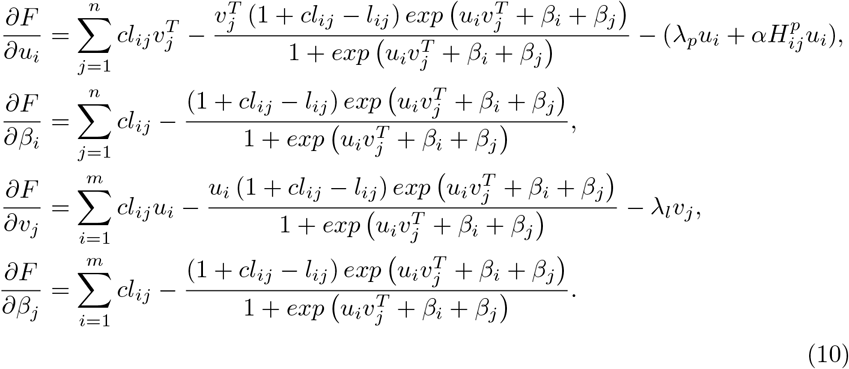

Once the latent matrices *U*, *V*, *β*_*i*_ and *β*_*j*_ are calculated, the presence probability of a protein *i* in a subcellular location can be estimated by the logistic function in formula 1. However for a new protein the latent factors *u* and *b* are not available. Hence, for a new protein the presence probability in subcellular location *j* is estimated as follows:

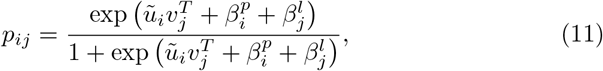

where 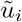 is the weighted average of the latent vectors of *k*_2_ nearest neighbors of *i*, as follows:

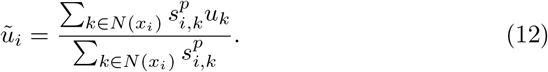

Eventually a threshold can be applied on probabilities to assign the subcellular locations to proteins.

### 2.2. Datasets and evaluation criteria

Evaluating the protein subcellular prediction methods is a challenging task. In one hand, the standalone version of state-of-the-art methods are not available and on the other hand, the protein databases are updated quickly. Hence, to achieve a fair evaluation and comparison we have employed the same datasets and evaluation criteria as used in previous studies [2, 5, 6]. These datasets are summarized in Table ??. The Hum-mploc3.0, the BaCelLo IDS animals[45], and the Höglund IDS[33] datasets consist of two non-overlapping subsets for training and testing purposes while for DBMloc we have performed 5-fold cross validation. The training set of Hum-mploc 3.0, HumB, is constructed from Swiss-Prot database release 2012 01 (January 2012) and consists of 3122 proteins of which 1023 proteins are labeled with more than one subcellular locations and the rest are single location proteins. Alongside HumB, HumT is used as the testing set to evaluate the method’s performance. HumT is also constructed from Swiss-Prot database release 2015 05 (May 2015 release) and consists of 379 proteins of which 120 proteins are labeled with more than one subcellular locations and the rest are single location proteins. Each protein in Hum-mploc 3.0 is assigned to at least one of 12 subcellular locations (Centrosome, Cytoplasm, Cytoskeleton, Endoplasmic reticulum, Endosome, Extracellular, Golgi apparatus, Lysosome, Mitochondrion, Nucleus, Peroxisome, and Plasma membrane).

The training set of BaCelLo IDS animals dataset is extracted from Swiss-Prot release 48 (September 2005 release) containing 2597 single label proteins, while the testing set consists of 576 single label proteins extracted from Swiss-Prot between relese 49 and 54 (February 2006 and July 2007 releases). Each protein in BaCelLo IDS animal dataset is assigned one of four subcellular locations (Cytoplasm, Mitochondrion, Nucleus, and Secreted).

In the Höglund IDS dataset, the training set contains 5959 single label proteins extracted from Swiss-Prot release 42 and includes nine subcellular locations (Nucleus, Cytoplasm, Mitochondrion, Endoplasmic reticulum, Golgi apparatus, Peroxisome, Plasma membrane, Extracellular space, Lysosome, and Vacuole) while the testing set contains 158 single label proteins extracted from Swiss-Prot release 55.3 including six subcellular locations (Endoplasmic reticulum, Golgi apparatus, Peroxisome, Plasma membrane, Extracellular space, and Lysosome). Accordingly, to train PSL-Recommender we only used 2682 proteins of training set that their subcellular location existed in the test set.

**Table 1:** Datasets summary

|  | Hum-mPLoc 3.0 |  | BaCelLo |  | Höglund |  | DBMloc |
| --- | --- | --- | --- | --- | --- | --- | --- |
|  | Train | Test | Train | Test | Train | Test | All |
| Proteins count | 3129 | 379 | 2597 | 576 | 5959 | 158 | 3056 |
| Labels count | 4229 | 541 | 2597 | 576 | 5959 | 158 | 6112 |
| Locations count | 12 |  | 4 |  | 6 |  | 6 |

Unlike the previous datasets, the DBMLoc dataset does not have a separate training and testing dataset. This dataset contains 3054 double locational proteins with paired subcellular locations: (cytoplasm and nucleus), (extracellular and plasma membrane), (cytoplasm and plasma membrane), (cytoplasm and mitochondrion), (nucleus and mitochondrion), (endoplasmic reticulum and extracellular) and (extracellular and nucleus). We have performed 5-fold cross validation technique to produce training and testing sets on this dataset.

We assessed PSL-Recommender performance against other methods by using customized ACC and F1 – mean over subcellular locations for evaluation of multi-label classification performance methods which is introduced by [46] and used by other state-of-the-art methods for this problem. ACC is the average of 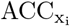 of all proteins in the test set, calculated for each protein as follows:

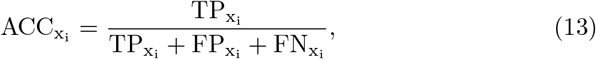

where, 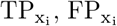 and 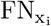 are number of true positive prediction, number of false positive predictions, and number of false negative predictions for protein *x*_*i*_, respectively.

The F1 – mean is the average of 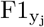 of all subcellular locations, where F1 of subcellular location *y*_*j*_ is the harmonic mean of 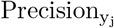 and 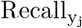, defined as follows:

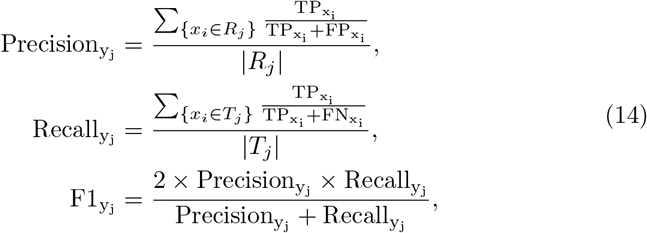

where, *R*_*j*_ and *T*_*j*_ are sets of predicted proteins for location *y*_*j*_ and true proteins for location *y*_*j*_, respectively.

Alongside, SherLoc2 [5] applied two other evaluation criterias, named ACC2 (ratio of correctly predicted proteins) and AVG (average fraction of called instances) which are defined as follow:

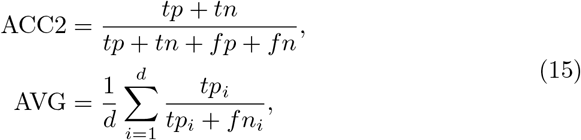

where, d denotes the number of subcellular locations and tp, tn, fp, and fn indicate the number of true positive, true negative, false positive, and false negative instances, respectively.

#### 2.2.1. Learning Hyperparameters

For all datasets, to prevent overfitting in tuning hyperparameters, they were learned from a comprehensive dataset, HumB, which is not considered in testing stage. It means that these hyperparameters are considered for all datasets. Since HumB -among the four mentioned datasets-contains both the single label and multi label PSL data, this dataset has been used for tuning task. Following 5-fold cross validation procedure is applied on HumB and hyperparameters were chosen empirically by maximizing the F1 – mean: HumB is devided into 5 equal subsets and PSL-Recommender is trained on union of 4 subsets and one other subset was hold for test the F1 – mean. This process is repeated 5 times, such that each time one of the 5 subsets is used as validation set and other 4 subsets are put together to form a training set.

For each set of hyperparameters, whole 5-fold process is repeated for 20 times and average of F1 – mean has been calculated. Due to the large search space, a grid-search procedure is applied for selecting the hyperparameters.

The weight of similarity measures used to build the protein similarity matrix was picked from 1 to 10 by step of 1. The dimension of latent space, *r*, was selected between 1 and the number of subcellular locations by step of 1. The weighting factor for positive observations, *c*, was chosen between 5 and 80 by step of 1. The number of nearest neighbors for constructing 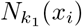 in equation 7, *k*_1_, was selected from 1 to 60 by step of 1. Similarly, The number of nearest neighbors for constructing 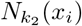, in equation 12, *k*_2_, was selected from 1 to 60 by step of 1. The variance controlling parameters, λ^*p*^ and λ_*l*_, were chosen form {2^−5^, 2^−4^, …, 2^1^}. Impact factor of nearest neighbors in equation 8, *α*, was picked from {2^−5^, 2^−4^, …, 2^2^}. Finally, The learning rate of the gradient descent criteria, *θ*, was selected from {2^−5^, 2^−4^, …, 2^0^}.

Table 2 represents the learned hyperparameters using HumB dataset. For all datasets, these learned hyperparameters are considered to construct the models.

**Table 2:** Learned hyperparameters based on HumB dataset. (*r* is latent space dimension, *c* is weighting factor for positive observations, *k*_1_ is the number of nearest neighbors for constructing 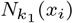, *k*_2_ is the number of nearest neighbors for constructing 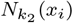, λ_*p*_ variance controlling parameterof proteins, λ_*l*_ variance controlling parameter of subcellular locations, *α* is the impact factor of nearest neighbors, and *θ* is the learning rate of the gradient descent criteria)

| | $r$ | $c$ | $k_1$ | $k_2$ | $\lambda_p$ | $\lambda_l$ | $\alpha$ | $\theta$ |
| --- | --- | --- | --- | --- | --- | --- | --- | --- |
| Value | 10 | 11 | 4 | 10 | 0.25 | 0.5 | 2 | 1 |

## 3. Results and discussion

PSL-Recommender can be employed to predict the subcellular protein localization in different species. Accordingly, we evaluated the performance of PSL-Recommender on different datasets and compared it to other state-of-the-arts methods. We further investigated the role of each protein similarity measures that are employed by the PSL-Recommender.

### 3.1. Comparison with the State-of-art method

We have first employed the Hum-mPLoc 3.0 [2] human protein dataset to compare the performance of PSL-Recommender to six methods that were introduced for protein localization in human. The methods include YLoc+ [6], iLoc-Hum [47], WegoLoc [48], mLASSO-Hum [49] and Hum-mPloc 3.0. The F1 – score for each location and the ACC and F1 – mean of all methods on Hum-mploc 3.0 dataset is depicted in Table 3.

**Table 3:** Comparison of PSL-Recommender on Human proteins dataset(Hum-mPloc 3.0) with other methods.

| Location | Yloc+ |  |  | iLoc-Human |  |  | WegoLoc |  |  | mLASSO-Hum |  |  | Hum-mPloc3.0 |  |  | PSL-Recommender |  |  |
| --- | --- | --- | --- | --- | --- | --- | --- | --- | --- | --- | --- | --- | --- | --- | --- | --- | --- | --- |
|  | pre | re | F1 | pre | re | F1 | pre | re | F1 | pre | re | F1 | pre | re | F1 | pre | re | F1 |
| Centrosome | - | - | - | 0 | 0 | 0 | 0.75 | 0.14 | 0.23 | 0.59 | 0.59 | 0.59 | 0.75 | 0.55 | <u>0.63</u> | 0.94 | 0.69 | **0.80** |
| Cytoplasm | 0.55 | 0.85 | 0.67 | 0.5 | 0.54 | 0.52 | 0.69 | 0.53 | 0.60 | 0.93 | 0.51 | 0.66 | 0.76 | 0.73 | <u>0.74</u> | 0.81 | 0.78 | **0.79** |
| Cytoskeleton | - | - | - | 0 | 0 | 0 | 0.32 | 0.34 | 0.33 | 0.9 | 0.22 | 0.35 | 0.8 | 0.68 | <u>0.74</u> | 0.97 | 0.70 | **0.82** |
| ER | 0.71 | 0.12 | 0.21 | 0 | 0 | 0 | 0.73 | 0.2 | 0.31 | 0.74 | 0.49 | <u>0.59</u> | 0.83 | 0.37 | 0.51 | 0.91 | 0.72 | **0.80** |
| Endosome | - | - | - | 0 | 0 | 0 | 0.25 | 0.07 | 0.11 | 0.38 | 0.2 | 0.26 | 0.58 | 0.47 | **0.52** | 0.63 | 0.31 | <u>0.41</u> |
| Extracellular | 0.39 | 0.85 | 0.54 | 0.62 | 0.62 | 0.62 | 0.67 | 0.77 | **0.71** | 0.16 | 0.69 | 0.26 | 0.5 | 0.46 | 0.48 | 0.66 | 0.71 | <u>0.68</u> |
| Golgi apparatus | 0.1 | 0.05 | 0.07 | 0.6 | 0.3 | 0.4 | 0.6 | 0.15 | 0.24 | 0.72 | 0.65 | <u>0.68</u> | 0.69 | 0.45 | 0.55 | 0.86 | 0.59 | **0.70** |
| Lysosome | 0 | 0 | 0 | 0.5 | 0.13 | 0.2 | 0.2 | 0.13 | 0.15 | 0.55 | 0.75 | 0.63 | 0.71 | 0.63 | <u>0.67</u> | 1 | 0.55 | **0.71** |
| Mitochondrion | 0.65 | 0.43 | 0.52 | 0.95 | 0.33 | 0.49 | 0.79 | 0.73 | 0.76 | 0.83 | 0.88 | <u>0.85</u> | 0.78 | 0.75 | 0.76 | 0.93 | 0.86 | **0.90** |
| Nucleus | 0.41 | 0.57 | 0.48 | 0.54 | 0.7 | 0.61 | 0.65 | 0.64 | 0.64 | 0.85 | 0.7 | <u>0.76</u> | 0.75 | 0.71 | 0.73 | 0.83 | 0.91 | **0.87** |
| Peroxisome | 0.07 | 0.5 | 0.13 | 1 | 0.5 | <u>0.67</u> | 0.5 | 1 | <u>0.67</u> | 0.29 | 1 | 0.44 | 1 | 1 | **1** | 1 | 1 | **1** |
| Plasma membrane | 0.41 | 0.44 | 0.42 | 0.42 | 0.33 | 0.37 | 0.44 | 0.53 | 0.48 | 0.58 | 0.56 | <u>0.57</u> | 0.65 | 0.44 | 0.52 | 0.77 | 0.73 | **0.75** |
| ACC | 0.45 |  |  | 0.41 |  |  | 0.50 |  |  | <u>0.65</u> |  |  | 0.63 |  |  | **0.77** |  |  |
| F1-mean | 0.34 |  |  | 0.32 |  |  | 0.44 |  |  | 0.56 |  |  | <u>0.65</u> |  |  | **0.77** |  |  |

As seen in Table 3, PSL-Recommender significantly outperforms the F1 – mean and ACC of all other methods improving the best method by 12% in both F1 – mean and ACC. Also, in 10 out of 12 subcellular locations, PSL-Recommender has the best performance amongst all methods while in the other two locations it has the second best performance. The most significant improvements have been observed in Centrosome, ER (Endoplasmic Reticulum) and Plasma Membrane showing 17%, 21% and 18% improvement respectively over the second best method.

It is only in Endosome that PSL-Recommender shows unsatisfactory results (41% F1 - score). This is while other methods also fail to provide good results for this location such that the best method (Hum-mPLOC 3.0) only achieves 52% F1 - score. Moreover, for Extracellular, WegoLoc slightly (3%) outperforms PSL-Recommender.

To show the performance of PSL-Recommender on other species we have employed previously introduced datasets that include proteins from animals and eukaryotes. We then compared the results to five state-of-the-art methods including [2, 6, 8, 33, 50]. The results are depicted in Table 4.

**Table 4:**
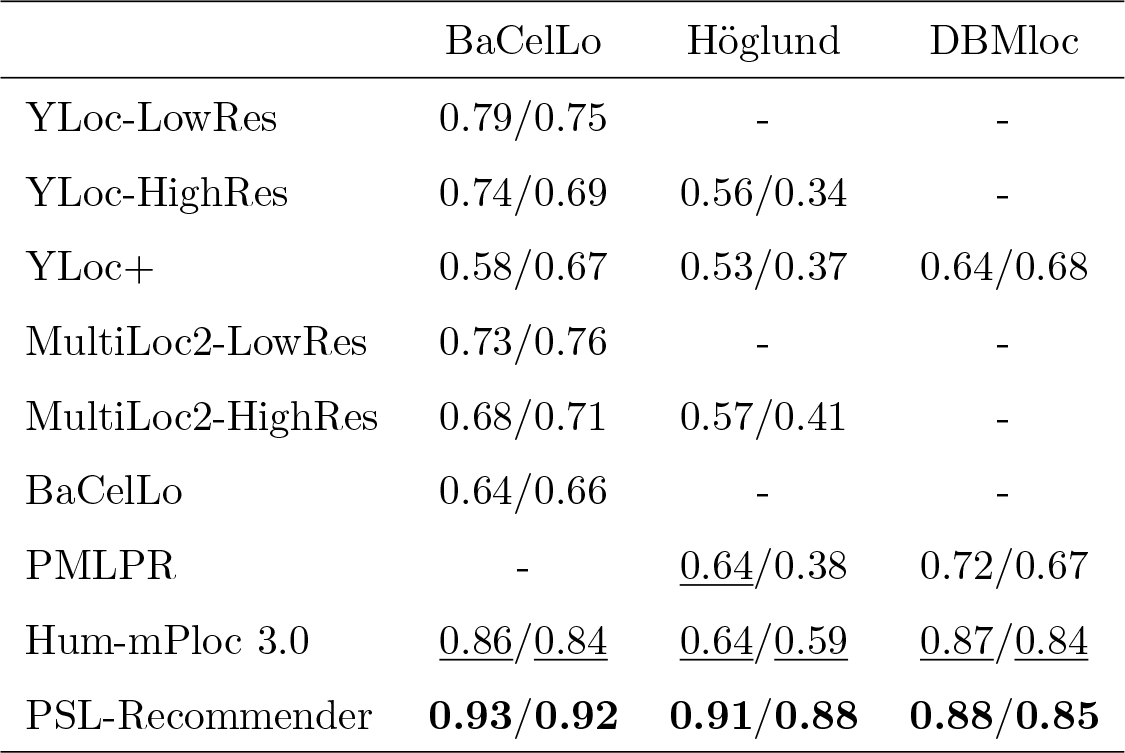
Comparison of PSL-Recommender ACC/F1 – mean on other species proteins datasets with state-of-the-art methods.

|  | BaCelLo | Höglund | DBMloc |
| --- | --- | --- | --- |
| YLoc-LowRes | 0.79/0.75 | - | - |
| YLoc-HighRes | 0.74/0.69 | 0.56/0.34 | - |
| YLoc+ | 0.58/0.67 | 0.53/0.37 | 0.64/0.68 |
| MultiLoc2-LowRes | 0.73/0.76 | - | - |
| MultiLoc2-HighRes | 0.68/0.71 | 0.57/0.41 | - |
| BaCelLo | 0.64/0.66 | - | - |
| PMLPR | - | <u>0.64</u>/0.38 | 0.72/0.67 |
| Hum-mPloc 3.0 | <u>0.86</u>/0.84 | <u>0.64</u>/0.59 | <u>0.87</u>/0.84 |
| PSL-Recommender | **0.93/0.92** | **0.91/0.88** | **0.88/0.85** |

As seen in Table 4, PSL-Recommender outperforms all methods in all datasets by both F1 – mean and ACC. In Höglund IDS animals dataset, PSL-Recommender significantly outperforms the second best method by 27% and 29% in F1 – mean and ACC respectively. In BaCelLo IDS animals dataset, the improvement over the second best method is 7% in F1 – mean and 8% in ACC, while in DBMloc dataset, PSL-Recommender slightly improves the second best method by 1% in both F1 – mean and ACC.

In order to compare PSL-Recommender performance with some other prominent works like SherLoc2 [5], WoLF PSORT [4], and Euk-mPloc [17], we have investigated AVG and ACC2 of PSL-Recommender results over two data set BaCelLo IDS animals and Höglund IDS animals and compared them with reported results in SherLoc2 paper which is represented in Table 5.

**Table 5:** Comparisons of PSL-RecommendeR, SherLoc2, MultiLoc2, WoLF PSORT, and Euk-mPloc performance with respect to AVG/ACC2.

| Method | BaCelLo | Höglund |
| --- | --- | --- |
| PSL-Recommender | **0.92/0.96** | **0.86/0.97** |
| SherLoc2 | <u>0.76/0.71</u> | <u>0.39/0.54</u> |
| MultiLoc2 | 0.75/0.68 | 0.38/0.57 |
| WoLF PSORT | 0.69/<u>0.71</u> | 0.24/0.56 |
| Euk-mPloc | 0.48/0.58 | 0.18/0.22 |

Table 6 is also demonstrate great performance of PSL-Recommender with re-spect to AVG and ACC2. PSL-Recommender shown great improvement by 16% in AVG and 25% in ACC2 over BaCelLo IDS dataset and also outperforming results over Höglund IDS dataset with 47% and 40% improvement over AVG and ACC2 respectively.

**Table 6:** PSL-Recommender ACC/F1 – mean comparisons by using different features.

| Features | BaCelLo | Höglund | DBMLoc | Hum-mPloc 3.0 |
| --- | --- | --- | --- | --- |
| PSSM | 0.69/0.53 | 0.63/0.26 | 0.81/0.77 | 0.33/0.17 |
| STRING | - | - | - | 0.44/0.40 |
| GO | 0.93/0.91 | 0.90/0.87 | 0.86/0.84 | 0.76/0.75 |
| PSSM+STRING | - | - | - | 0.46/0.37 |
| GO +PSSM | 0.93/0.92 | 0.91/0.88 | 0.88/0.85 | 0.77/0.76 |
| GO+STRING | - | - | - | 0.78/0.77 |
| All | 0.93/0.92 | 0.91/0.88 | 0.88/0.85 | 0.77/0.77 |

It also worth mentioning that, for PSL prediction problem, to the best of our knowledge, PMLPR [8] is the only recommender system based method that employs the well-known network-based inference(NBI) [51] approach. As seen in Table 4, PSL-Recommender outperforms PMLPR by 50% and 18% in F1 – mean, and also 27% and 16% in ACC on Höglund and DBMloc datasets, repectively.

### 3.2. Impact of each similarity matrix

The proteins similarity matrix is used for neighborhood regularization and also the prediction step. To acquire this matrix PSL-Recommender combines three sources of protein similarity measures (PSSM similarity, String-DB interactions similarity and GO terms semantic similarity) using weighted averaging. The weights are acquired through the learning process.

To investigate the impact of different similarity measures, we repeated previous experiments using different combination of similarity measures. Table 6. shows the result of each combination on all datasets. As can be seen in Table 6., those combinations excluding the GO terms semantic similarities do not provide reliable predictions showing that GO terms semantic similarities play an important role in protein subcellular localization.

It should be noted that GO terms are not available for all proteins. In the absence of GO terms semantic similarities, PLS-Recommender is still able to provide acceptable results for DBMLoc and BacelLo datasets but its performance significantly drops for Höglund and Hum-mPloc 3.0.

Moreover, the usage of String protein-protein interaction scores is only limited to datasets that contain proteins from single species. Since DBMLoc, BacelLo, and Höglund datasets contain proteins from multiple species we were unable to use String interaction scores in these datasets.

### 3.3. Stability of the PSL-Recommender

Choosing appropriate hyperparameters plays a vital role in the performance of a model. As mentioned in section 2.2.1, the models for all of the datasets constructed by same set of hyperparameters based on HumB dataset (Table 2). The results of using these hyperparameters represented on Table 3, Table 4, and Table 5 for all datasets.

In order to investigate the stability of the models, for each datasets, the hyperparametrs are selected according to their training set by applying 5-fold cross validation with similar procedure which is explained in section 2.2.1 By considering different hyperparameters, F1 – mean reached to 0.92, 0.90, and 0.89 and ACC get to 0.94, 0.92, and 0.89 for BaCelLo IDS, Höglund IDS, and DBMloc, respectively.

For each datasets, by applying selected hyperparameters with respect to their training set, the F1 – mean and ACC can be increased only by 2 percent. It can be concluded that, despite large number of hyperparameters PSL-Recommender is a stable method for PSL prediction.

## 4. Conculusion

In the absence of efficient experimental methods, computational tools play an important role for predicting protein subcellular localizations. Yet, there is still much room for improving the prediction accuracy of these methods. In this paper, we introduced PSL-Recommender, a recommender system that employs logistic matrix factorization for efficient prediction of protein subcellular localization. By evaluating on human and animals datasets it was shown that PSL-Recommender significantly outperforms other state-of-the-art methods. However, we believe that the performance of PSL-Recommender can be improved further by employing a better approach for searching the parameter space. The standalone version of PSL-Recommender and all the datasets are available online at: https://github.com/RJamali/PSL-Recommender

## Acknowledgements

This work is supported by Iran National Science Foundation.

